## Supplemental Materials for "A versatile information retrieval framework for evaluating profile strength and similarity"

#### *Information retrieval tasks for profile evaluation*

| **Task** | **Query group** | **Reference group** |
|---|---|---|
| *Phenotypic activity* | Replicate profiles of a perturbation | Replicate profiles of controls |
| *Phenotypic distinctiveness* | Replicate profiles of a perturbation | Replicate profiles of other perturbations |
| *Phenotypic consistency* | Perturbations that share the same annotation | Perturbation that do not have the same annotation |

**Supplementary Table 1**. Information retrieval tasks for profile evaluation.

#### *Effect and replicability in phenotypic activity assessment*

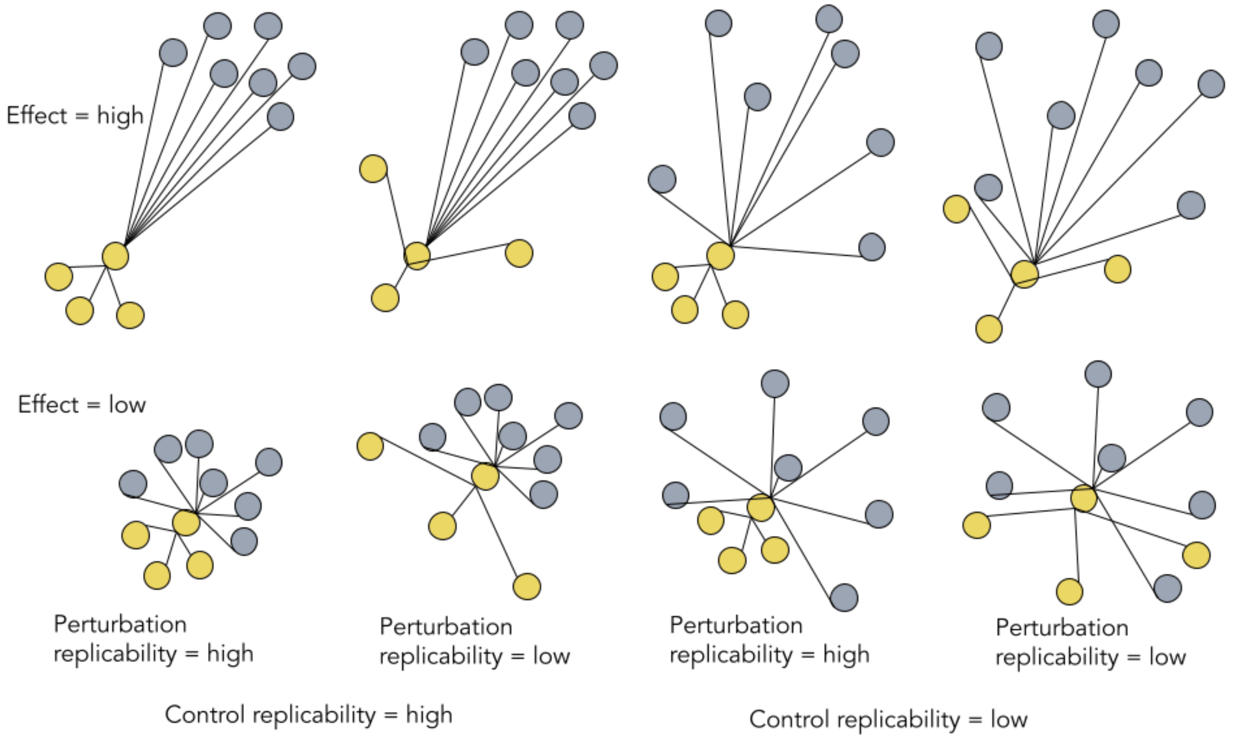


**Figure S1.** Schematic illustration of effect and replicability in phenotypic activity of a perturbation. Depending on effect strength induced by a perturbation and the amount of technical replicability variation, it may be nearly impossible to distinguish between effect and replicability.

#### *Additional comparisons of mAP with alternatives on simulated data*

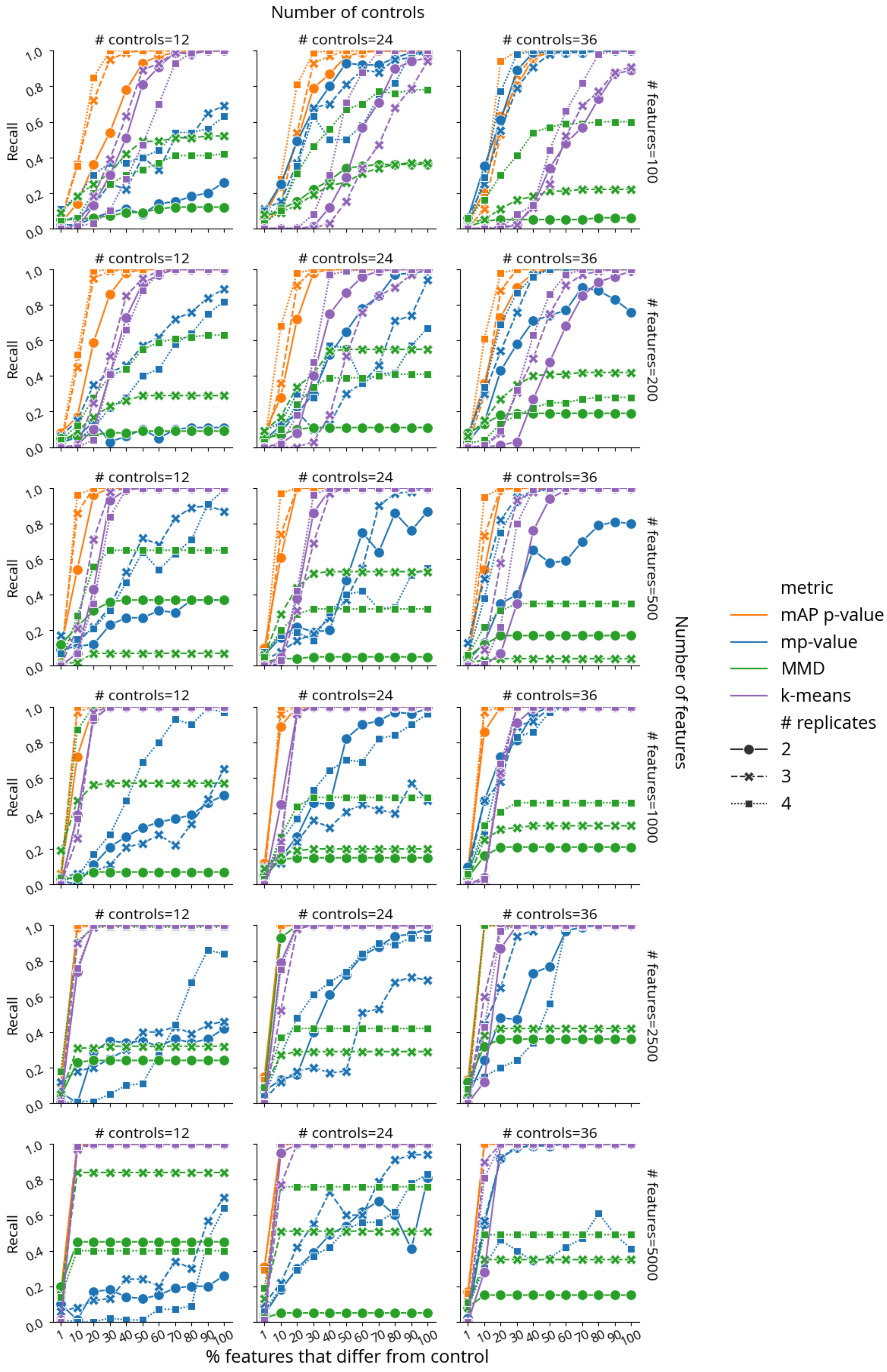

Number of controls
### controls=12
### controls=24
### controls=36
Recall
### features=100
### features=200
### features=500
### features=1000
### features=2500
### features=5000
Number of features
% features that differ from control
metric
mAP p-value
mp-value
MMD
k-means
### replicates
2
3
4

**Figure S2.** Benchmarking retrieval performance of mAP p-value (orange), mp-value (blue), MMD p-value (green), and k-means clustering (purple) for assessing phenotypic activity on simulated data, where unperturbed and perturbed features are sampled from $\mathcal{N}(0,1)$ and $\mathcal{N}(1,1)$, correspondingly. Recall indicates the percentage of 100 simulated perturbations under each condition that were called accurately by each method (as distinguishable from negative controls, or not). The horizontal axis probes what proportion of the features in the profile was different from controls. Marker and line styles indicate different numbers of replicates per perturbation (# replicates of 2, 3, and 4). Columns correspond to the different number of controls (# controls of 12, 24, and 36). Rows correspond to different profile sizes (# features being 100, 200, 500, 1000, 2500, and 5000).

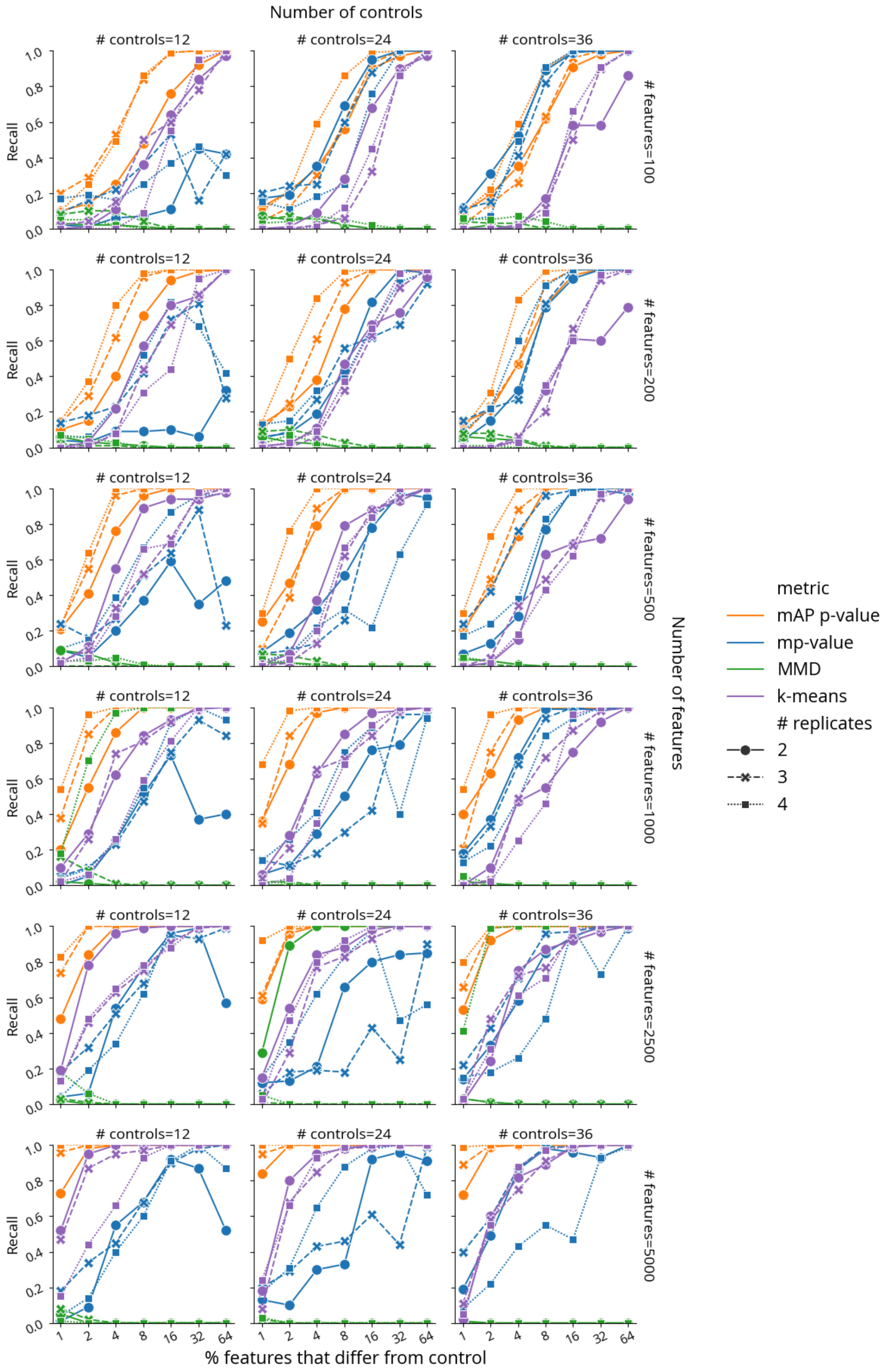

Number of controls
### controls=12
### controls=24
### controls=36
Recall
### features=100
### features=200
### features=500
### features=1000
### features=2500
### features=5000
Number of features
% features that differ from control
metric
mAP p-value
mp-value
MMD
k-means
### replicates
2
3
4

**Figure S3.** Benchmarking retrieval performance of mAP p-value (orange), mp-value (blue), and MMD p-value for phenotypic activity on simulated data, where unperturbed and perturbed features are sampled from $\mathcal{N}(0,1)$ and $\mathcal{N}(2,2)$, correspondingly. Recall indicates the percentage of 100 simulated perturbations under each condition that were called accurately by each method (as distinguishable from negative controls, or not). The horizontal axis probes what proportion of the features in the profile was different from controls. Marker and line styles indicate different numbers of replicates per perturbation (# replicates of 2, 3, and 4). Columns correspond to the different number of controls (# controls of 12, 24, and 36). Rows correspond to different profile sizes (# features being 100, 200, 500, 1000, 2500, and 5000).

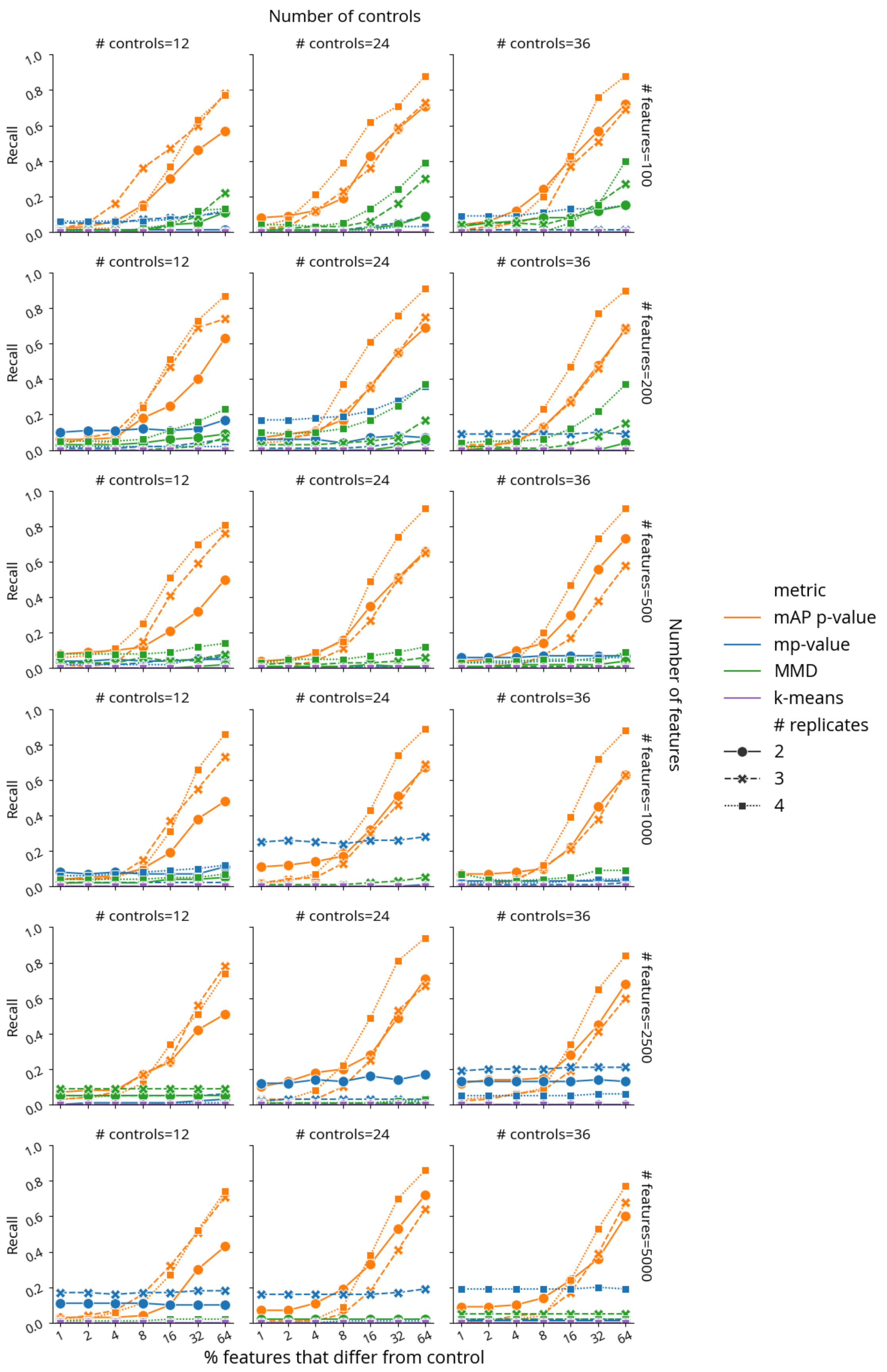

Number of controls
### controls=12
### controls=24
### controls=36
Recall
1.0
0.8
0.6
0.4
0.2
0.0
### features=100
### features=200
### features=500
### features=1000
### features=2500
### features=5000
Number of features
metric
mAP p-value
mp-value
MMD
k-means
### replicates
2
3
4
1
2
4
8
16
32
64
% features that differ from control

**Figure S4.** Benchmarking retrieval performance of mAP p-value (orange), mp-value (blue), and MMD p-value for phenotypic activity on simulated data, where unperturbed and perturbed features are sampled from Cauchy(0,1) and Cauchy(10,1), correspondingly. Recall indicates the percentage of 100 simulated perturbations under each condition that were called accurately by each method (as distinguishable from negative controls, or not). The horizontal axis probes what proportion of the features in the profile was different from controls. Marker and line styles indicate different numbers of replicates per perturbation (# replicates of 2, 3, and 4). Columns correspond to the different number of controls (# controls of 12, 24, and 36). Rows correspond to different profile sizes (# features being 100, 200, 500, 1000, 2500, and 5000).

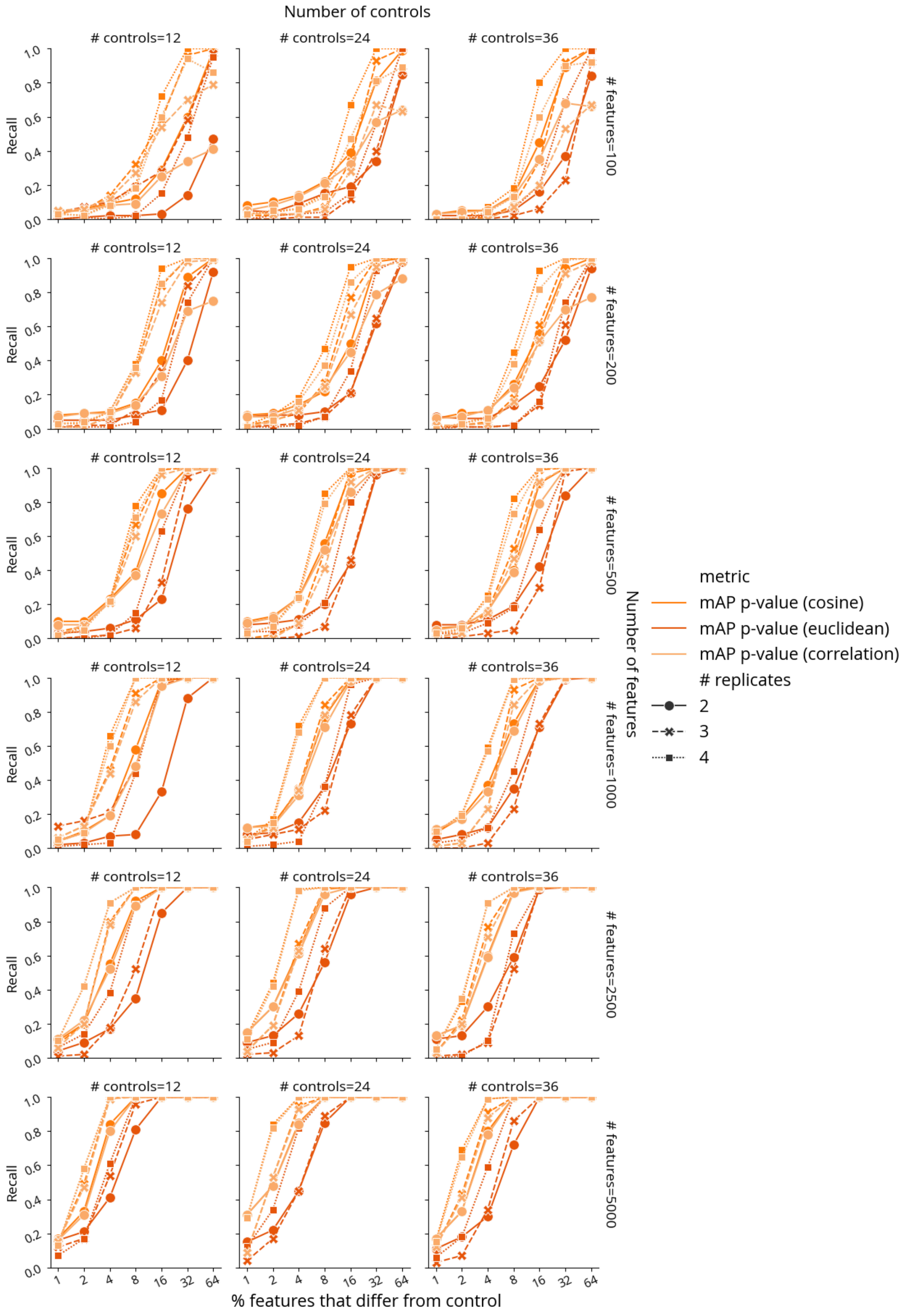

Number of controls
### controls=12
### controls=24
### controls=36
Recall
### features=100
### features=200
### features=500
### features=1000
### features=2500
### features=5000
Number of features
% features that differ from control
metric
mAP p-value (cosine)
mAP p-value (euclidean)
mAP p-value (correlation)
### replicates
2
3
4

**Figure S5.** Benchmarking retrieval performance of mAP p-value calculated using different distance metrics. We compared mAP p-value retrieval with the default choice of cosine distance (orange) against mAP p-value with Euclidean distance (dark orange) and correlation (light orange) for phenotypic activity on simulated data, where unperturbed and perturbed features are sampled from $\mathcal{N}(0,1)$ and $\mathcal{N}(1,1)$, correspondingly. Recall indicates the percentage of 100 simulated perturbations under each condition that were called accurately by each method (as distinguishable from negative controls, or not). The horizontal axis probes what proportion of the features in the profile was different from controls. Marker and line styles indicate different numbers of replicates per perturbation (# replicates of 2, 3, and 4). Columns correspond to the different number of controls (# controls of 12, 24, and 36). Rows correspond to different profile sizes (# features being 100, 200, 500, 1000, 2500, and 5000).

#### *Additional mAP applications to the Cell Health dataset*

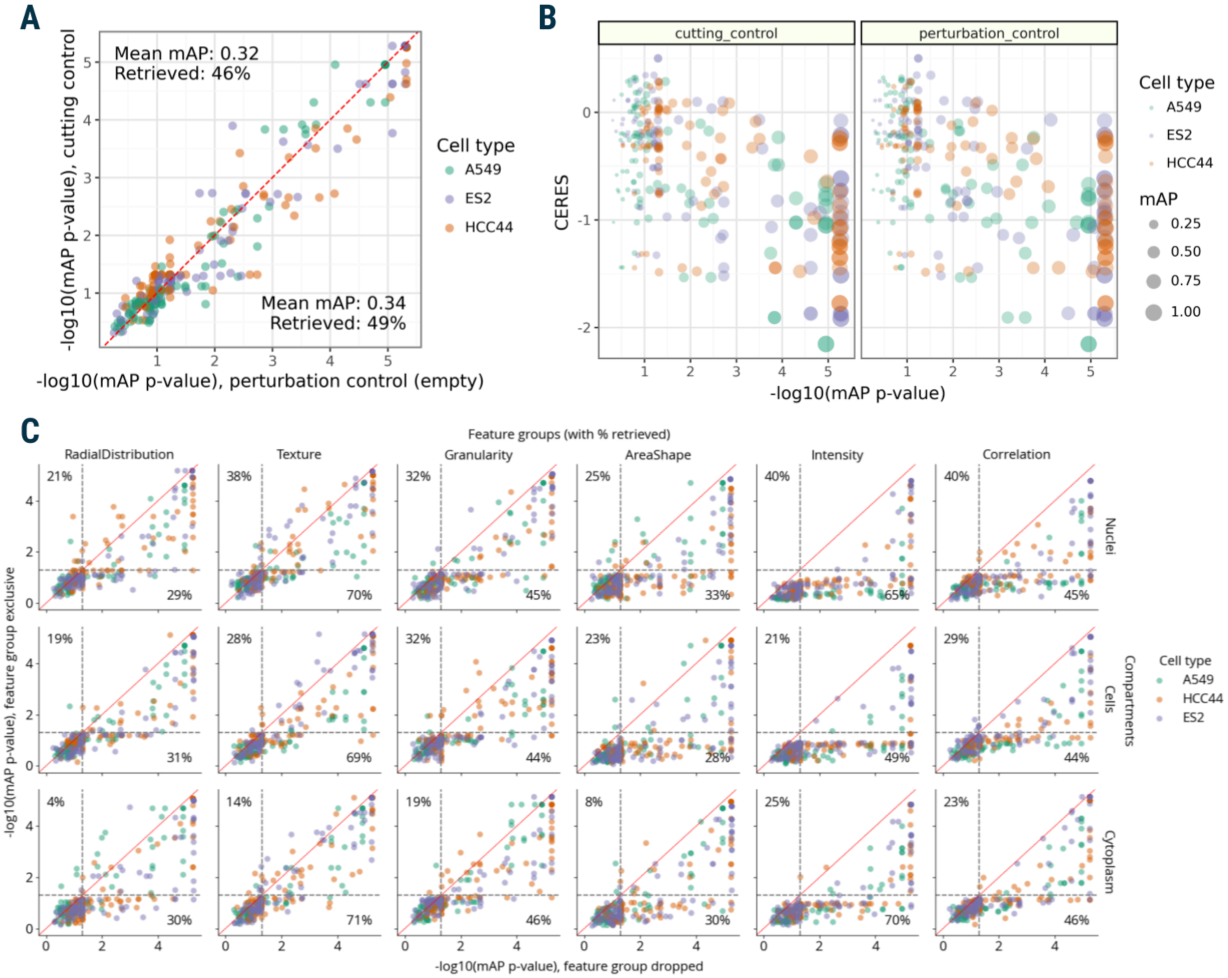


**Figure S6**. Additional results for the mAP framework application to morphological profiling using Cell Health data. **A**: Correlation between mAP *p*-values that assessed the phenotypic activity of perturbations by guide replicate retrievability against non-targeting cutting controls and perturbation controls (empty wells). **B**: Correlation between mAP *p*-values that assessed the phenotypic activity of perturbations by guide replicate retrievability against non-targeting cutting controls and perturbation controls (empty wells). **C**: mAP *p*-values calculated to assess influence of individual fluorescent channels on guide phenotypic activity against controls by either dropping a channel or including only that single channel (percent retrieved is shown for each axis), these results can be compared to 49% retrieved when all channels' data is available (on average across all three cell lines, as in **B**). **D**: mAP is calculated to assess the phenotypic consistency of guides annotated with related target genes (against guides annotated with other genes) in three cell lines individually.

#### *mAP application to cpg0004 and cpg0016[orf] datasets*

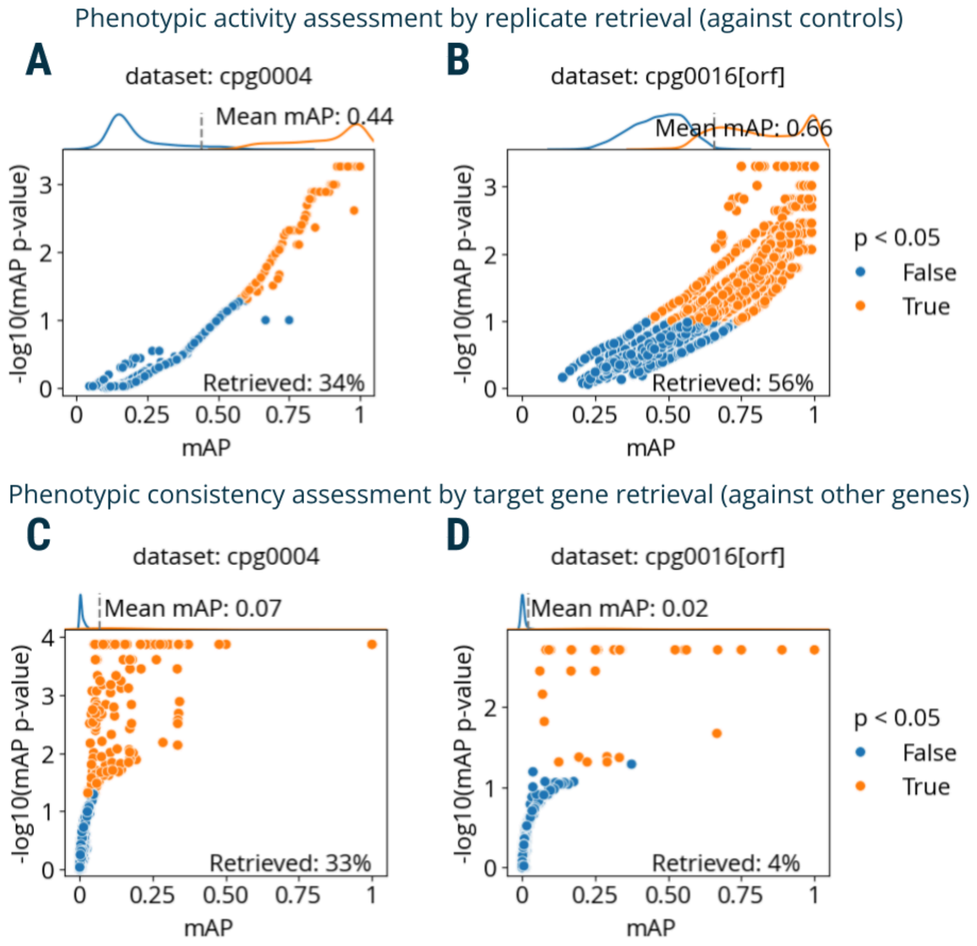


**Figure S7.** Additional results for the mAP framework application to morphological profiling of small molecule (cpg0004) and gene overexpression (cpg0016[orf]) perturbation datasets. **A**: mAP is calculated to assess the phenotypic activity of compounds by replicate retrievability against controls in cpg0004 data. **B**: mAP is calculated to assess the phenotypic activity of gene overexpression reagents by replicate retrievability against controls in cpg0016[orf] data. **C**: mAP is calculated to assess the phenotypic consistency of compounds annotated with related target genes (against compounds annotated with other genes) in cpg0004 data. **D**: mAP is calculated to assess the phenotypic consistency of gene overexpression reagents annotated with related protein complexes (against compounds annotated with other protein complexes) in cpg0016[orf] data.

#### *Additional results for AP application to the Perturb-seq dataset*

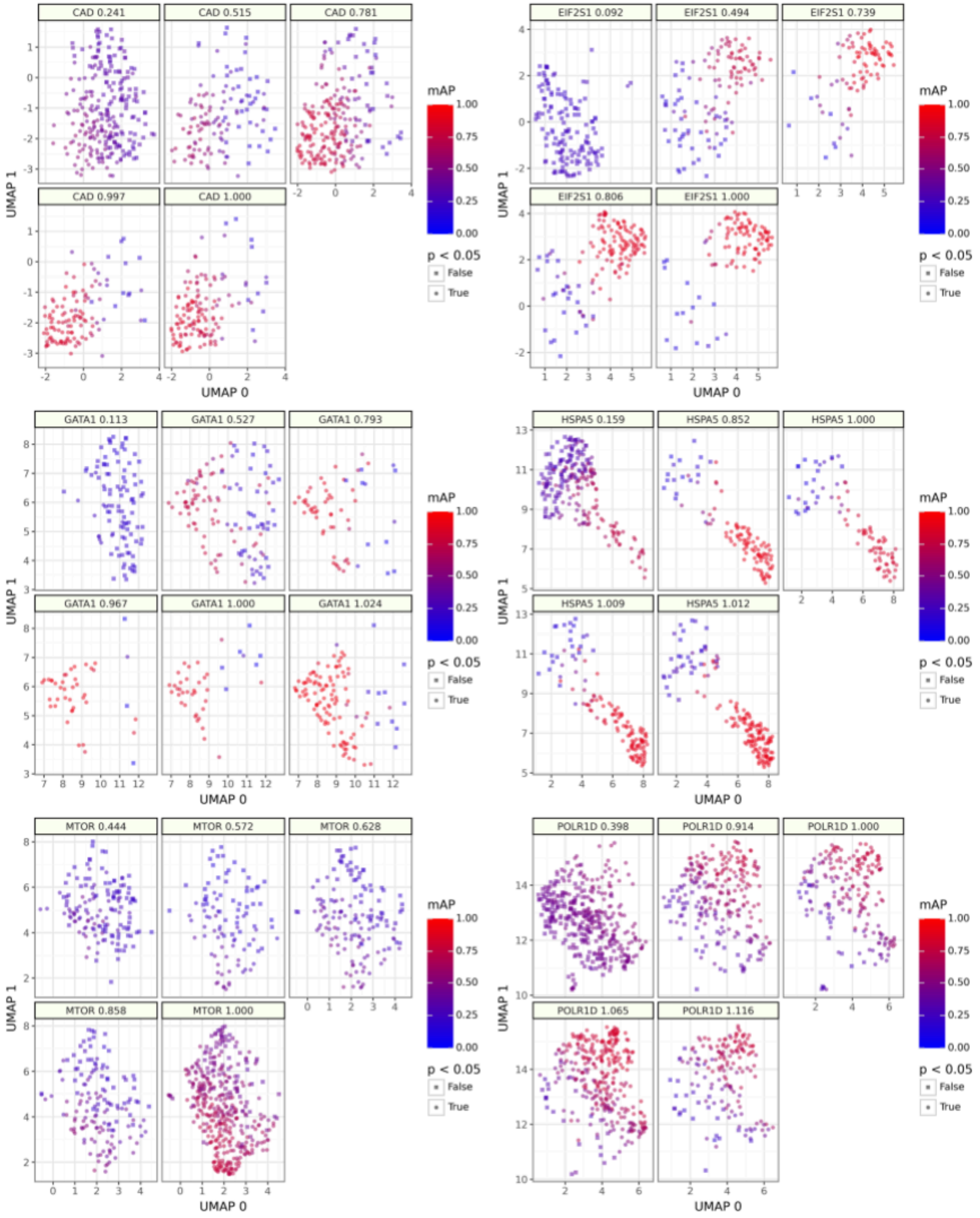

**Figure S8.** Single-cell profile AP scores assessing phenotypic activity for individual guides with varying mismatches relative to the perfect-matching guide (a subset of data shown in Figure 5A). Guides with higher relative activity scores demonstrate higher single-cell AP values. UMAP embeddings were created using n_neighbors=30 and cosine similarity as a distance measure.

#### *Additional mAP applications to the Mitocheck dataset*

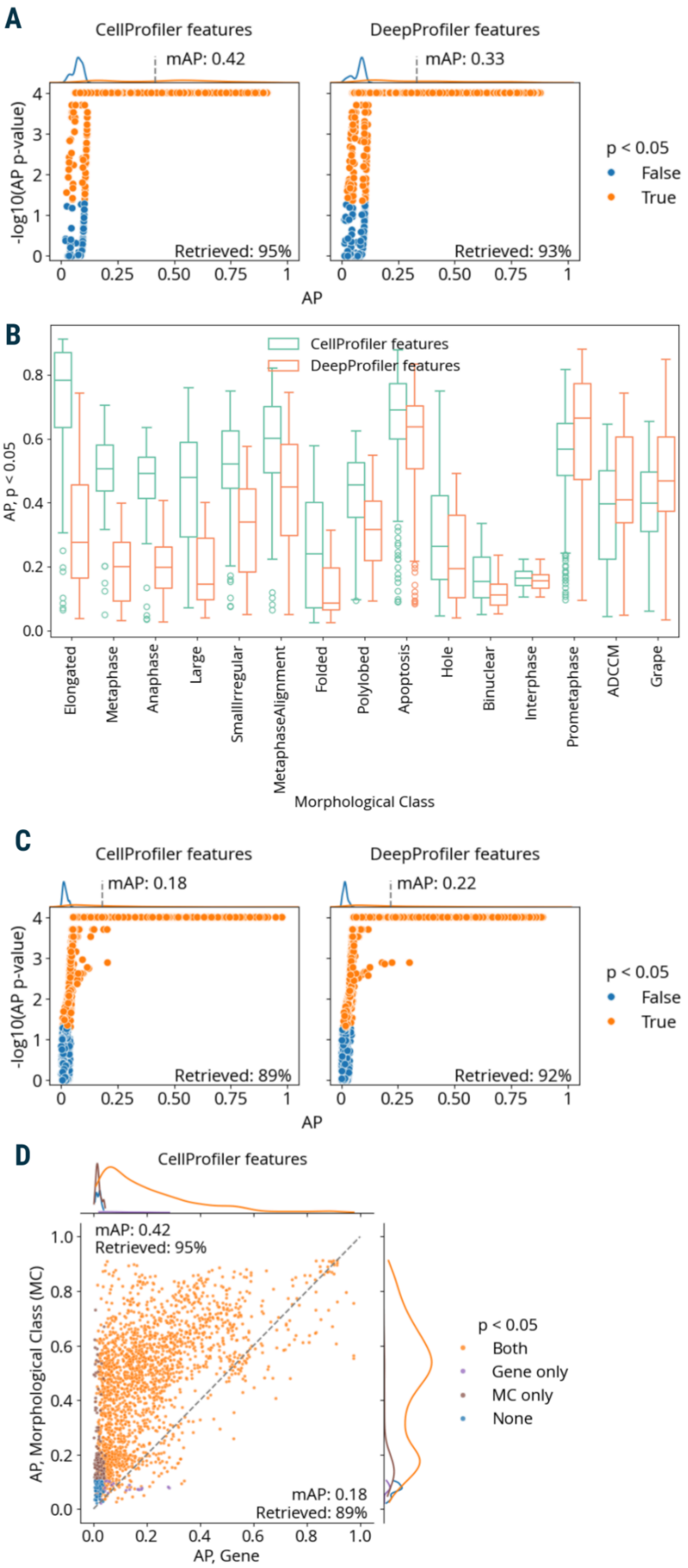
A
CellProfiler features
DeepProfiler features
mAP: 0.42
mAP: 0.33
-log10(AP p-value)
Retrieved: 95%
Retrieved: 93%
AP
p < 0.05
False
True
B
CellProfiler features
DeepProfiler features
AP, p < 0.05
Elongated
Metaphase
Anaphase
Large
SmallIrregular
MetaphaseAlignment
Folded
Polylobed
Apoptosis
Hole
Binuclear
Interphase
Prometaphase
ADCCM
Grape
Morphological Class
C
CellProfiler features
DeepProfiler features
mAP: 0.18
mAP: 0.22
-log10(AP p-value)
Retrieved: 89%
Retrieved: 92%
AP
p < 0.05
False
True
D
CellProfiler features
mAP: 0.42
Retrieved: 95%
mAP: 0.18
Retrieved: 89%
AP, Morphological Class (MC)
AP, Gene
p < 0.05
Both
Gene only
MC only
None

**Figure S9.** Additional results for the mAP framework application to single-cell profiling using Mitocheck morphological data. **A**: AP scores can be used to evaluate the power of CellProfiler and DeepProfiler features to classify morphological classes in Mitocheck data. AP scores capture the ability to retrieve single cells annotated with the same morphological class against negative controls. **B**: A subset of AP scores with $p < 0.05$ from panel A is shown to evaluate the power of CellProfiler and DeepProfiler features to classify specific morphological classes. **C**: AP scores can be used to evaluate the power of CellProfiler and DeepProfiler features to classify target genes in Mitocheck data. AP scores capture the ability to retrieve single cells annotated with the same target gene against negative controls. **D**: A subset of AP scores obtained using CellProfiler features from panels **A** and **C** is shown to compare retrieval of morphological classes and target genes.

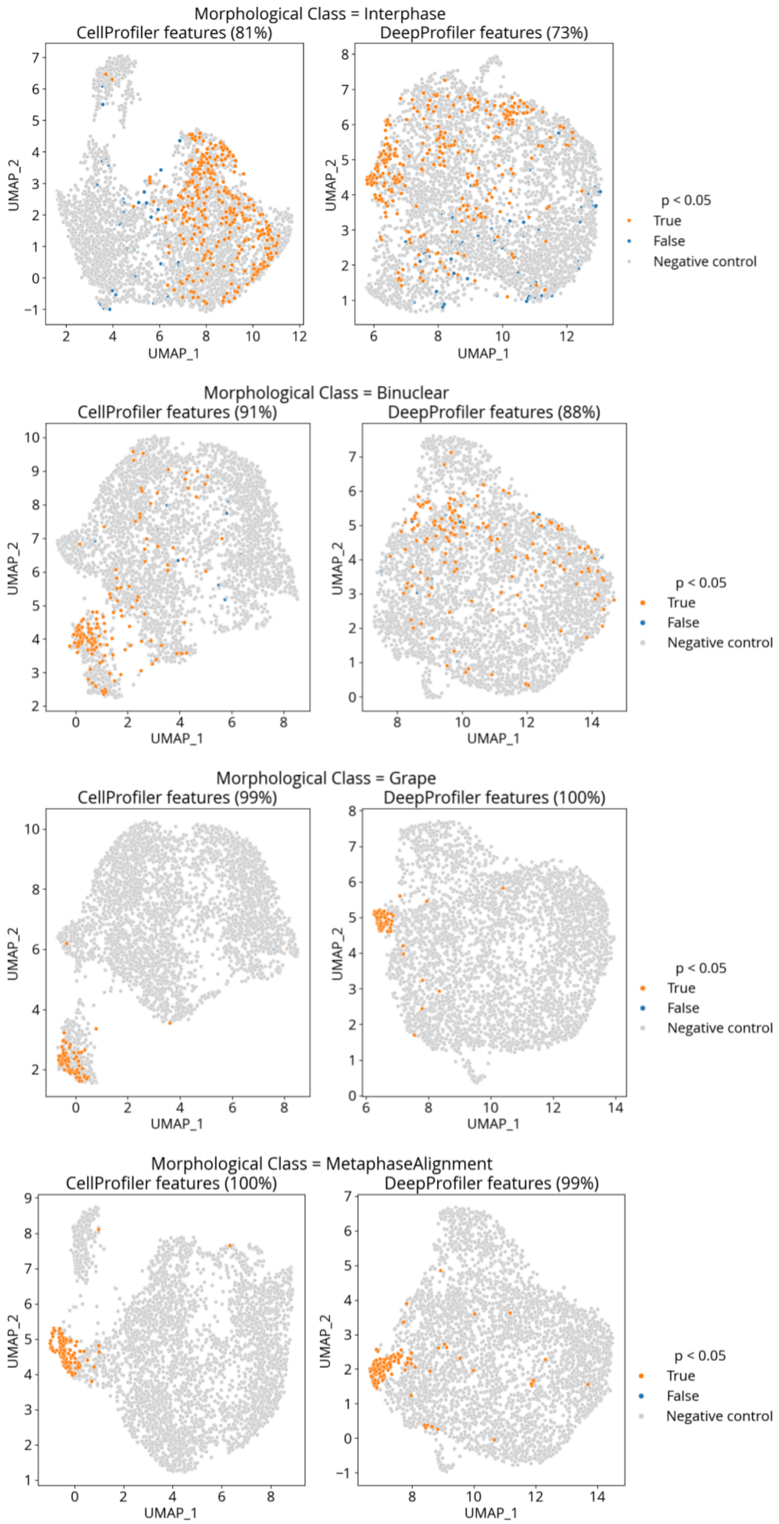


**Figure S10.** Single-cell profile AP scores assessing phenotypic activity for exemplar morphological classes in Mitocheck data with CellProfiler and DeepProfiler features (a subset of data shown in Figure 5C). Classes with higher retrieval rates (Grape, MetaphaseAlignment) demonstrate tighter clustering compared to those with lower rates (Interphase, Binuclear with DeepProfiler features). UMAP embeddings were created using n_neighbors=30 and cosine similarity as a distance measure.

#### *Runtime analysis*

We optimized the mAP framework implementation for efficient processing of large datasets. We analyzed runtime performance for phenotypic activity assessment (retrieval of each perturbation replicates against negative controls) across all datasets used in this study.

Calculating mAP involves three steps:

1. Profile grouping

The first step involves using metadata to match profiles that form a query group (i.e., replicate profiles of each perturbation) and a reference group (i.e., replicate profiles of a negative control). Because mAP is calculated only based on query-to-control and within-the-query-group distances, we can avoid computing the full distance matrix, which in the case of large datasets is not computationally feasible. This also enables flexible profile grouping by various block designs, which can be useful for reducing computational complexity or accounting for confounding variables. Our implementation achieves efficient matching through structured sampling of profile pairs based on predefined constraints on column values.

Approximate overall time complexity can be estimated as following:

- Best case (`sameby` constraints limit pairs effectively): $O(np)$, where $n$ is the number of rows and $p$ is the number of `diffby` constraints.
- Worst case (if `sameby` constraints are loose, allowing many pairs): $O(n^2p)$.

2. mAP calculation

At the second step, mAP score per query group (perturbation) is calculated as the mean of individual AP scores calculated per each query (replicate), as described in *Methods: mAP calculation* section of the paper. Main steps here include pairwise distance calculation for positive (within query group) and negative (query to reference group) profile pairs and distance sorting, while AP score calculation cost is negligible to those. Time complexity of these steps can be estimated as following:

- Best case (many small groups): $O(nmd)+O(nm \times log(nm))$, where $d$ is profile dimensionality and $m$ is the average profile group size.
- Worst case (few large groups): $O(n^2d)+O(n^2 \times log(n^2))$.

In our implementation, distance computation uses multithreading to speed up calculations.

3. *p*-value calculation

Permutation testing for mAP score includes calculating unique group sizes and constructing corresponding null distributions, as described in the *Methods: Assigning significance to mAP scores section* of the paper. Approximate time complexity can be estimated as following:

- Best case (few unique group sizes, small null size): $O(n \times log(n))$
- Worst case (many unique group sizes, large null size): $O(mcs)$, where $c$ is the number of unique group sizes and $s$ is the null size.

Random AP score calculations to construct null distributions are also implemented using multithreading. Null distributions for the specific group sizes are cached locally and reused.

Overall, time complexity depends on the number of perturbations of the dataset, the number of metadata constraints for profile grouping, sizes of perturbation groups (the number of perturbation replicates) and control groups (the number of control replicates), profile dimensionality (the number of features in a profile), and selected null size. Choosing the null size can be challenging as it requires balancing test resolution (an ability to differentiate between significance levels) and compute required to provide adequate sampling coverage given a particular dataset.

We report runtime across datasets for phenotypic activity assessment (retrieval of perturbation replicates against negative controls) in **Supplementary Table 2**. Calculations were performed using AMD Ryzen Threadripper PRO 7995WXSx192 CPU and 769 GB 2,133 MHz DDR4 RDIMM memory. We report p-value calculation runtime with cleared cache; in practice this only occurs when running the analysis for the first time and the cost becomes negligible after that.

| **Dataset** | **# perts** | **# reps** | **# ctrls** | **Null size** | **Runtime, sec** | | | |
|---|---|---|---|---|---|---|---|---|
| | | | | | **Profile grouping** | **AP calculation** | **p-value calculation** | **Total** |
| Cell Health (HCC44) | 100 | 6 | 192 | $1\times10^6$ | 0.03 | 0.22 | 2.1 | 2.3 |
| cpg0004 | 1558 | 5 | 24 | $5\times10^4$ | 0.08 | 0.56 | 1.55 | 2.2 |
| cpg0016[orf] | 13739 | 4 | 16 | $2\times10^4$ | 0.11 | 2.32 | 0.02 | 2.5 |
| nELISA (Cell Painting) | 300 | 4 | 64 | $1\times10^5$ | 0.04 | 0.53 | 0.3 | 0.9 |
| Perturb-seq (guide-level, gene retrieval) | 25 | 5-6 | 10 | $1\times10^4$ | 0.02 | 0.06 | 0.005 | 0.08 |
| Perturb-seq (single-cell, guide retrieval) | 128 | 32-561 | 472 | $5\times10^5$ | 0.66 | 4.78 | 108.39 | 114 |
| Mitocheck (single-cell, Morphological class retrieval, CellProfiler) | 15 | 36-409 | 3900 | $5\times10^5$ | 0.2 | 5.19 | 252.3 | 179.9 |
| Mitocheck (single-cell, Gene retrieval, CellProfiler) | 59 | 3-153 | 3900 | $5\times10^5$ | 0.22 | 5.26 | 889.1 | 894.6 |

**Supplementary Table 2**. Runtime across datasets for phenotypic activity assessment.
